# Carbon source, cell density, and the microbial community control inhibition of *V. cholerae* surface colonization by environmental nitrate

**DOI:** 10.1101/2024.12.31.630902

**Authors:** Jamaurie James, Renato ERS Santos, Paula I Watnick

## Abstract

The intestinal diarrheal pathogen *Vibrio cholerae* colonizes the host terminal ileum, a microaerophilic, glucose-poor, nitrate-rich environment. In this environment, *V. cholerae* respires nitrate and increases transport and utilization of alternative carbon sources via the cAMP receptor protein (CRP), a transcription factor that is active during glucose scarcity. Here we show that *V. cholerae* nitrate respiration in aerated cultures is under control of CRP and, therefore, glucose availability. *V. cholerae* nitrate respiration results in extracellular accumulation of nitrite because *V. cholerae* does not possess the machinery for nitrite reduction. This nitrite inhibits *V. cholerae* biofilm formation via an as yet unelucidated mechanism that depends on the high cell density master regulator HapR. The genome of *Paracoccus aminovorans*, an intestinal microbe shown to enhance *V. cholerae* biofilm accumulation in the neonatal mouse gut and predispose household contacts to cholera, encodes enzymes that reduce nitrite to nitrogen gas. We report that, in nitrate-supplemented co-cultures, *P. aminovorans* metabolizes the nitrite generated by *V. cholerae* and, thereby, enhances *V. cholerae* surface accumulation. We propose that *V. cholerae* biofilm formation in the host intestine is limited by nitrite production but can be rescued by intestinal microbes such as *P. aminovorans* that have the capacity to metabolize nitrite. Such microbes increase *V. cholerae* colonization of the host ileum and predispose to infection.

**Importance:** *V. cholerae* colonizes the terminal ileum where both oxygen and nitrate are available as terminal electron acceptors. *V. cholerae* biofilm formation is inhibited by nitrate due to its conversion to nitrite during *V. cholerae* respiration. When co-cultured with a microbe that can further reduce nitrite, *V. cholerae* surface accumulation in the presence of nitrate is rescued. The contribution of biofilm formation to ileal colonization depends on the composition of the microbiota. We propose that the intestinal microbiota predisposes mammalian hosts to cholera by consuming the nitrite generated by *V. cholerae* in the terminal ileum. Differences in the intestinal abundance of nitrite-reducing microbes may partially explain the differential susceptibility of humans to cholera and the resistance of non-human mammalian models to intestinal colonization with *V. cholerae*.

## Introduction

The Gram-negative pathogen *Vibrio cholerae* is a facultative anaerobe that colonizes the ileum to cause the acute and severe diarrheal disease cholera (1). In an infant mouse model, the principal *V. cholerae* attachment factor is the type IV toxin co-regulated pilus (2). In contrast, on environmental surfaces, *V. cholerae* constructs a three-dimensional surface attached structure known as the VPS- dependent biofilm by synthesizing a VPS exopolysaccharide scaffold via enzymes encoded by the *vps* genes as well as the three secreted lectins RbmA, RbmC, and Bap1 (3–6). After secretion, these lectins bind to the VPS exopolysaccharide and form bridges between neighboring cells and between cells and the surface to erect a stable, three-dimensional structure. There is evidence from infant mouse studies that this VPS-dependent biofilm increases small bowel colonization *in vivo* when members of the human intestinal microbiota that predispose to cholera are present (7–10).

*V. cholerae* is found in the small and large bowel of the infant mouse during infection, but only in the large bowel of germ-free or streptomycin-treated adult mice (11, 12). Large bowel colonization is asymptomatic in mice and independent of the toxin co-regulated pilus, a principal colonization factor (13). Asymptomatic carriage of *V. cholerae* also occurs in humans and may reflect large bowel colonization (14, 15). This suggests that the conditions encountered by *V. cholerae* in the small intestine are essential for activation of virulence determinants and development of disease. Availability of the terminal electron acceptors oxygen and nitrate varies along the length of the intestine, and these modulate virulence behaviors in pathogens other than *V. cholerae* (16–22). The partial pressure of oxygen is highest in the duodenum at approximately 60 mmHg and decreases to under 10 mmHg in the ileum (22). Despite the lower oxygen concentrations, *V. cholerae* colonization of the small bowel relies heavily on aerobic respiration that utilizes the terminal oxidases cytochrome bd-type ubiquinol oxidase I (bd-1) and the heme copper oxidase Cbb3 (23, 24).

Nitrate, which is generated through the actions of iNOS and NOX1, is present in the adult mouse ileum at concentrations approaching 6 mM and at much lower concentrations in other regions of the small intestine (20, 25).

*V. cholerae*’s use of nitrate as a terminal electron acceptor is limited. While other bacterial genomes encode multiple systems for nitrate reduction to nitrite, the *V. cholerae* genome harbors only a periplasmic nitrate reductase encoded by the *napDABC* genes (24, 26). Other bacteria are able to further reduce nitrite to nitrogen gas via a pathway comprised of multiple reductive enzymes that carry out the following reactions: nitrite (NO_2_ ^-^)→nitric oxide (NO)→nitrous oxide (N2O)→nitrogen gas (N2) (27) In contrast, *V. cholerae* reduces nitrate to nitrite but cannot carry out additional reductions (28). This results in accumulation of conditionally toxic nitrite in the extracellular environment (29).

The *V. cholerae* genes required for nitrate respiration under hypoxic laboratory conditions have been exhaustively identified by Tnseq and include the *nap* genes, the *nqr* genes, which encode the Na+- translocating NADH-quinone reductase, and the *ccm* cytochrome maturation genes (30). These authors found a small but significant colonization defect for a Δ*napA* mutant in a streptomycin-treated adult mouse model, although colonization was quantified in the caecum and colon only (29, 30). A microarray analysis of *V. cholerae* in human cholera stool found that the *nap* genes were highly expressed as compared with laboratory conditions (31). However, expression varied greatly between the three patients studied, suggesting significant effects of the intestinal environment.

We hypothesized that nitrate reduction might play an important role in *V. cholerae* colonization of the terminal ileum. Because this environment contains oxygen, we began by defining regulation of *V. cholerae* nitrate reduction in aerobic cultures and its effect on surface colonization. Here we describe an unusual role for two transcription factors, the cAMP receptor protein (CRP) and HapR in regulation of *V. cholerae* nitrate reduction to nitrite and inhibition of *V. cholerae* biofilm formation by nitrite, respectively.

CRP is a conserved global regulator of carbon metabolism that binds DNA in the presence of the second messenger cyclic adenosine monophosphate (cAMP) (32). cAMP is synthesized by the enzyme adenylate cyclase. When glucose is available, adenylate cyclase is inactive, cAMP is not synthesized, and CRP does not activate transcription. Therefore, the behavior of a Δ*crp* mutant should approximate that of wild-type (WT) *V. cholerae* in a glucose-rich environment. When glucose is scarce as is the case in Luria-Bertani (LB) broth, adenylate cyclase is active. The resulting elevated levels of intracellular of cAMP lead to formation of a cAMP-CRP complex, which activates transcription of hundreds of genes including those encoding transporters and enzymes required to utilize alternative sugars and amino acids as carbon sources (33–35). *V. cholerae* CRP also represses many processes required for virulence including cholera toxin, the toxin co-regulated pilus, and biofilm formation (33–38). In general, genes in the CRP regulon change their transcription in response to the availability of glucose as a carbon source.

The *V. cholerae* transcription factor HapR regulates transcription at high cell density. At low cell density, *hapR* mRNA is destabilized by the four small RNA’s *qrr1-4* (39). Therefore, Δ*hapR* mutant behavior should approximate that of WT *V. cholerae* at low cell density. As the cell density increases, *qrr*’s are repressed in response to four secreted *V. cholerae*-derived small molecules known as autoinducers, and *hapR* mRNA is stable and transcribed (40). CRP and HapR interact at many levels to regulate gene transcription. For instance, *hapR* transcription is activated by CRP (34, 41). In addition, CRP and HapR share many binding sites, and joint binding alters transcription of target genes (42).

We investigated the observation that *V. cholerae* biofilm formation is inhibited in the presence of nitrate. We find that nitrite accumulates extracellularly when *V. cholerae* is cultured aerobically in LB broth supplemented with nitrate. The accumulated nitrite, rather than nitrate, directly inhibits *V. cholerae* surface accumulation. Interestingly, CRP is essential for nitrite production in shaking but not static cultures, and this is manifested as resistance of the Δ*crp* mutant to inhibition of biofilm formation by environmental nitrate. We provide evidence that nitrite generation is inhibited by environmental oxygen and that the Δ*crp* mutant does not sufficiently deplete oxygen in aerated cultures to initiate nitrate reduction. We further report that a Δ*hapR* mutant is not susceptible to inhibition of *V. cholerae* biofilm accumulation by nitrite, but this is rescued by deletion of *crp*. While RNAseq analysis identified many genes and putative small RNAs that respond to nitrate in WT *V. cholerae* but not the Δ*hapR* mutant, none had a previously described role in biofilm formation or dispersal. This suggests that nitrite inhibits *V. cholerae* biofilm accumulation in glucose-poor environments only at high cell density by an as yet unelucidated transcriptional or post-transcriptional mechanism.

The *V. cholerae* machinery for oxygen and nitrate respiration is expressed under microaerophilic conditions, and nitrate is abundant in the ileal lumen (23). Therefore, we hypothesized that inhibition of *V. cholerae* biofilm accumulation by the nitrite resulting from nitrate reduction might be responsible for the minimal contribution of the VPS-dependent biofilm to *V. cholerae* colonization of the infant mouse small intestine (43, 44). *Paracoccus aminovorans* is a member of the microbiome of cholera- predisposed individuals and increases *V. cholerae* surface attachment in both laboratory conditions and the infant mouse (7, 8). Here we show that *P. aminovorans* respires the nitrite generated by *V. cholerae* in co-culture, thus rescuing *V. cholerae* biofilm accumulation. We propose that, upon arrival in the nitrate-rich terminal ileum, the contribution of VPS-dependent biofilm accumulation to *V. cholerae* intestinal colonization is determined by glucose availability via CRP, cell density via HapR, and the microbial environment via nitrite-reducing microbes. Thus, differences in nitrate and nitrite utilization by the intestinal microbial communities of neonatal and adult mammalian models as well as the human population may contribute to the observed variability in susceptibility to cholera.

## Results

### Nitrate inhibition of *V. cholerae* biofilm accumulation in shaking cultures is dependent on the transcription factor CRP

During ongoing studies of CRP, a subtle, qualitative difference was observed between the biofilms formed by wild-type (WT) *V. cholerae* and a Δ*crp* mutant in shaking but not static cultures. We hypothesized that this could be the result of differential availability of oxygen as a terminal electron acceptor in shaking WT *V. cholerae* and the Δ*crp* mutant cultures. To test this, we formed WT *V. cholerae* and Δ*crp* mutant biofilms in shaking LB cultures either alone or supplemented with 5 mM sodium nitrate or sodium fumarate, both of which can serve as alternative terminal electron acceptors. As shown in Figure 1A and B, nitrate significantly decreased biofilm accumulation by WT *V. cholerae* but not the Δ*crp* mutant. In contrast, fumarate did not alter WT *V. cholerae* biofilm accumulation, while it decreased biofilm accumulation by the *Δcrp* mutant. Nitrate and fumarate did not alter total growth (Fig 1B). This demonstrates that the impact of nitrate on *V. cholerae* biofilm formation is not simply a reflection of its ability to serve as an alternative electron acceptor. Because nitrate is present in the terminal ileum and, therefore, could be important for *V. cholerae* intestinal colonization, we elected to further investigate inhibition of WT *V. cholerae* biofilm formation by nitrate and the role played by CRP in this process.

**Figure 1:**
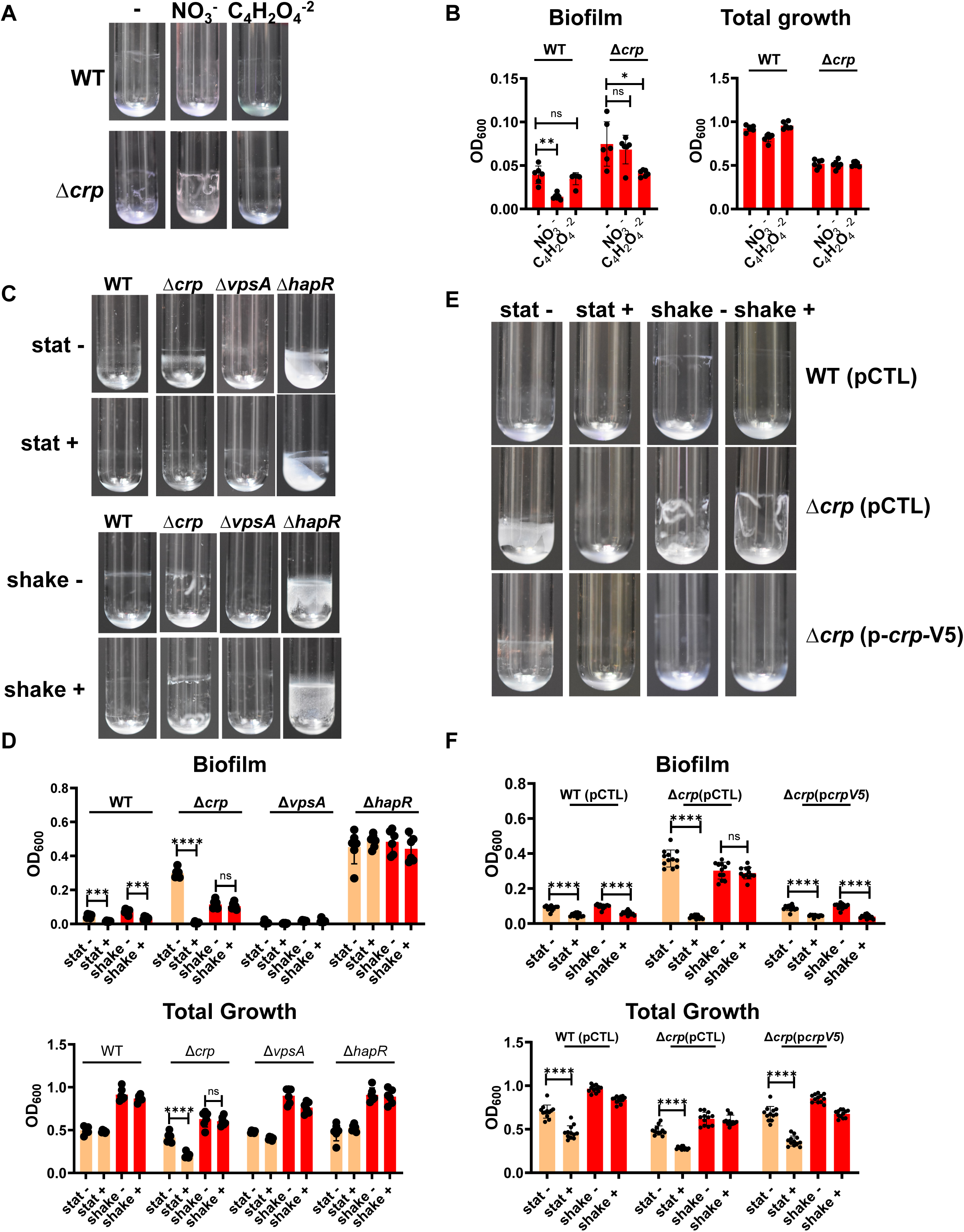
Inhibition of biofilm accumulation by nitrate is dependent on HapR in both aerated and static cultures and dependent on CRP in aerated cultures only. (A) Representative biofilm images and (B) quantification of biofilm formation and total growth by WT *V. cholerae* and a Δ*crp* mutant after approximately 18 hours of incubation at 27°C in LB alone (-) or LB supplemented with 5 mM sodium nitrate (NO_3_^-^) or 5 mM sodium fumarate (C_4_H_2_O ^-2^). Significance for total growth measurements was calculated using an ordinary one-way ANOVA with Dunnett’s multiple comparisons test. Significance for biofilm accumulation was calculated using a Brown-Forsythe and Welch’s ANOVA with Dunnett’s T3 multiple comparisons test. (C) Representative biofilm images and (D) quantification of biofilm formation and total growth by WT *V. cholerae* and the indicated mutants cultured under static (stat) or aerated (shake) conditions in LB alone (-) or supplemented with 5 mM NaNO_3_ (+). An expanded view of the WT biofilm quantification is shown in Fig S1A. (E) Representative biofilm images and (F) quantification of biofilm formation and total growth by WT *V. cholerae* and the indicated mutants. Strains were rescued with the empty vector pFLAG-CTC (pCTL) or the same vector encoding CRP (p*crp*-V5) as noted. Strains carrying plasmids were cultured with added ampicillin (100 µg/ml) and 0.5 mM IPTG. Significance for biofilm accumulation was calculated using a Welch’s t test. Significance for total growth measurements was calculated using a student’s t test. In all biofilm experiments, the mean of six biological replicates is shown, and error bars reflect the standard deviation. **** p<0.0001, *** p<0.001, ** p<0.01, * p<0.05, ns not significant.

The impact of nitrate on *V. cholerae* growth and surface association has principally been studied under hypoxic conditions (30, 45). To better understand the effect of nitrate on *V. cholerae* biofilm accumulation when both oxygen and nitrate are available as terminal electron acceptors, we quantified static and shaking biofilms formed by WT *V. cholerae*, a Δ*crp* mutant, and a Δ*vpsA* mutant in the presence and absence of nitrate. Nitrate supplementation caused a significant reduction in WT *V. cholerae* biofilm accumulation under both static and aerated conditions but did not alter total growth (Figs 1C and D, and S1A). In contrast, Δ*crp* mutant biofilm accumulation was unaffected by nitrate in shaking cultures but greatly reduced in static ones. Nitrate supplementation also reduced total growth of the Δ*crp* mutant in static cultures. However, the effect of nitrate on biofilm accumulation by the Δ*crp* mutant was approximately two orders of magnitude greater than that on its growth. Therefore, the reduction in total growth does not account for the decrease in Δ*crp* mutant biofilm accumulation. As expected, the Δ*vpsA* mutant accumulated very little biofilm under any of the conditions tested, and no difference in total growth was observed.

To establish that the inhibition of *V. cholerae* biofilm accumulation by nitrate was dependent on CRP in aerated cultures, we rescued the Δ*crp* mutant with a wild-type copy of *crp* provided on a plasmid behind an IPTG-inducible promoter. Inhibition of biofilm accumulation by nitrate in shaking cultures was restored to the Δ*crp* mutant by provision of *crp* in trans (Fig 1E and F). In all strains carrying the plasmid, small but significant decreases in total growth were observed only in static but not aerated cultures supplemented with nitrate, but this again did not account for the larger decreases in biofilm accumulation (Fig 1F). This supports the conclusion that CRP is essential for inhibition of biofilm accumulation by nitrate in shaking cultures. Since CRP is active only in glucose-poor environments, we propose that the available carbon source plays a role in inhibition of *V. cholerae* biofilm accumulation by nitrate in aerated cultures.

### A Δ*hapR* mutant is resistant to inhibition of biofilm accumulation by nitrate in both static and shaking cultures

CRP activates expression of HapR, the master regulator of high cell density quorum sensing, to repress biofilm formation (41, 46, 47). Furthermore, HapR is essential for biofilm dispersal at high cell density (48). To determine whether HapR plays a role in inhibition of biofilm accumulation by nitrate, we tested the impact of nitrate on biofilm accumulation by a Δ*hapR* mutant. Δ*hapR* mutant biofilm accumulation as well as total growth was unaffected by nitrate regardless of aeration (Fig 1C and D). Because a Δ*hapR* mutant is locked in the low cell density state, this result suggests that *V. cholerae* biofilm accumulation is resistant to nitrate at low cell density.

### Nitrate conversion to nitrite is essential for inhibition of WT and Δ*crp* mutant *V. cholerae* biofilm accumulation by nitrate

Because V*. cholerae* cannot reduce nitrate beyond nitrite, nitrite accumulates extracellularly when *V. cholerae* uses nitrate as an alternative electron acceptor. We reasoned, therefore, that inhibition of *V. cholerae* biofilm accumulation in the presence of nitrate could be a direct response to nitrate or a response to its metabolite nitrite. To test this, we formed WT, Δ*crp*, Δ*vpsA*, and Δ*hapR* mutant biofilms in LB supplemented either with 5 mM nitrate or nitrite under static and shaking conditions. As shown in Figs 2A and B, although aeration improved growth of all strains to some extent, WT *V. cholerae* biofilm accumulation in the presence of nitrate and nitrite was negligible independent of aeration, while the Δ*hapR* mutant formed robust biofilms. In contrast, while nitrate inhibited Δ*crp* mutant biofilm accumulation only in static cultures, nitrite inhibited Δ*crp* mutant biofilm accumulation regardless of aeration. Because the phenotypes of the Δ*crp* and Δ*hapR* mutants were distinct, we conclude that CRP and HapR act at least in part independently to modulate the *V. cholerae* response to nitrate.

**Figure 2:**
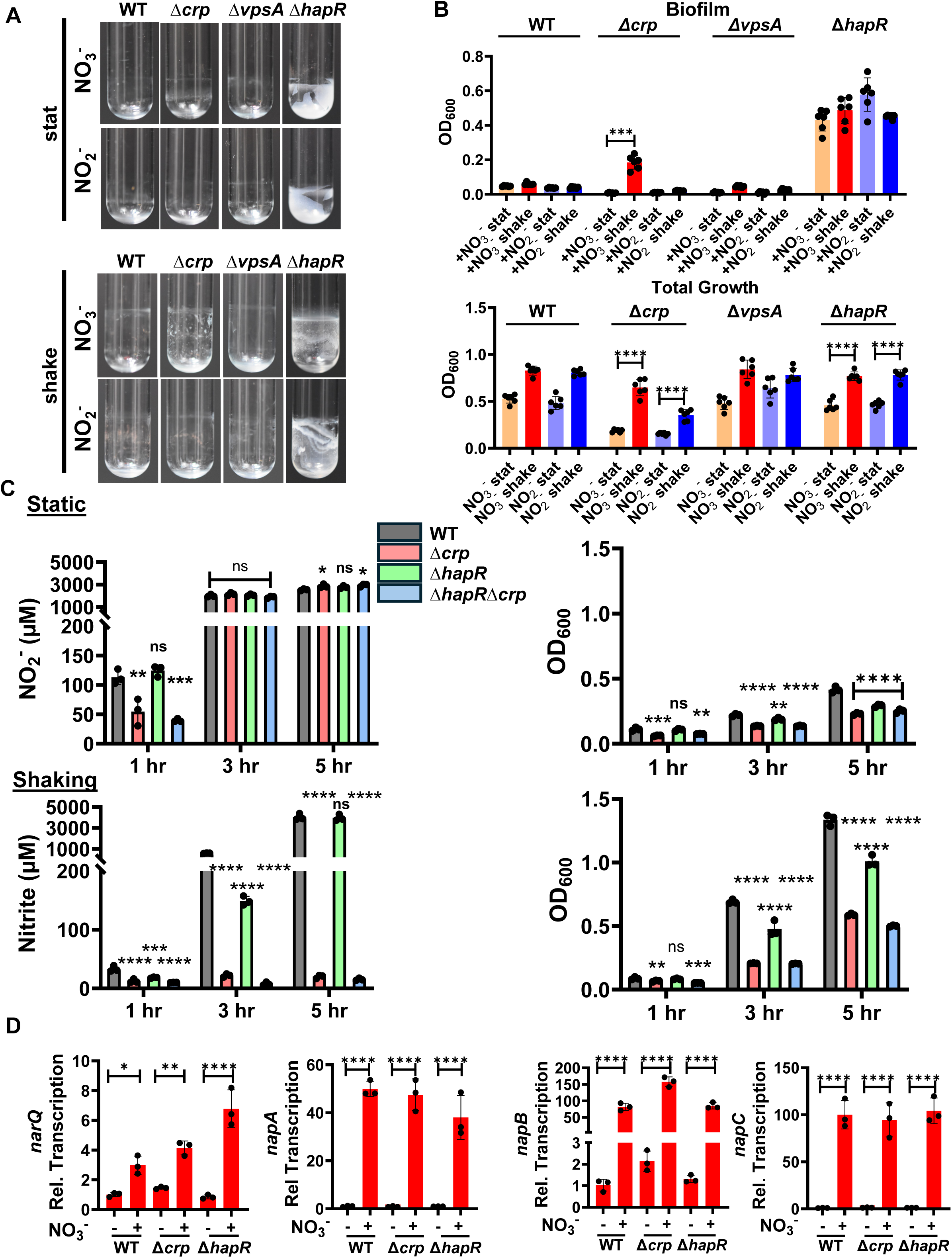
*V. cholerae* reduction of nitrate to nitrite is required for inhibition of biofilm accumulation and occurs in aerobic cultures only when CRP is present. (A) Representative images of biofilms formed by WT *V. cholerae* and the indicated mutants after approximately 20 hours of incubation at 27°C in LB supplemented with 5 mM nitrate (+NO_3_^-^) or 5 mM nitrite (+ NO_2_^-^). Biofilms were either formed statically (stat) or with aeration (shake). (B) Quantification of total growth and biofilm formation by the indicated strains cultured under the conditions indicated. The mean of six biological replicates is shown. Error bars reflect the standard deviation. For biofilm accumulation, significance was calculated using a Welch’s t test. For total growth, significance was calculated using a student’s t test. (C) Quantification of nitrite (NO_2_^-^) generation (left graph) and growth as assessed by OD_600_ (right graph) over time for the indicated strains cultured in LB supplemented with nitrate. Measurements were made under both static and shaking conditions. The mean of biological triplicates is shown. Error bars reflect the standard deviation. For both nitrite and growth measurements, significant differences at each time point were assessed using an ordinary one-way ANOVA with Dunnett’s multiple comparisons test. (D) qRT-PCR quantification of expression of the *narQ* and *nap* genes required for nitrite generation from nitrate in the indicated strains. Cultures were aerated and harvested at 8 hours. The mean of biological triplicates is shown. Error bars reflect the standard deviation. Significance was calculated using an ordinary one-way ANOVA with Dunnett’s multiple comparisons test. **** p<0.0001, *** p<0.001, ** p<0.01, * p<0.05, ns not significant.

Based on our findings, we hypothesized that nitrite is the direct inhibitor of *V. cholerae* biofilm accumulation at high cell density and that CRP is essential for efficient reduction of nitrate to nitrite in aerated cultures. To test this, we monitored growth and nitrite generation in static and aerated cultures of WT *V. cholerae* as well as the Δ*crp* and Δ*hapR* mutants in LB supplemented with nitrate. In aerated cultures of WT *V. cholerae*, nitrite generation from nitrate was delayed by 2-3 hours as compared with static cultures even though aeration improved growth (S1B). This suggests that oxygen availability inhibits nitrate reduction.

In static cultures, nitrite generation was similar for all strains tested despite significant differences in growth (Fig 2C). In contrast, in aerated conditions, the Δ*crp* mutant culture generated a negligible amount of nitrite over the course of the experiment, while the WT and Δ*hapR* mutant strains had converted almost all the nitrate to nitrite after 5 hours. While the Δ*crp* mutant grew more slowly than WT *V. cholerae*, this difference was too small to account for its inability to generate nitrite in aerated cultures. We hypothesize that oxygen inhibits nitrate generation by *V. cholerae* and that a Δ*crp* mutant, which does not initiate nitrate reduction to nitrite when cultured aerobically, may not be able to deplete aerated cultures of oxygen.

To further establish the role of nitrite in *V. cholerae* biofilm inhibition, we studied a Δ*napC* mutant that cannot generate nitrite. We first compared nitrite generation by WT *V. cholerae* and a Δ*napC* mutant in LB supplemented with nitrate. Nitrite concentrations in 6- and 24-hour Δ*napC* mutant cultures were not significantly different from that of sterile nitrate supplemented media (Fig S1C). We then tested the ability of the Δ*napC* mutant to form a biofilm in the presence of nitrate (Fig S1D and E). While nitrate supplementation inhibited biofilm accumulation by WT *V. cholerae*, it had no effect on the Δ*napC* mutant biofilm and no effect on total growth by either strain. We conclude that nitrate inhibits WT *V. cholerae* biofilm accumulation through conversion to nitrite.

### Differential transcription of the genes encoding NarQ and NAP complex components does not underlie the absence of nitrite generation by the *Δcrp* mutant in shaking cultures

Our results showed that CRP is essential for nitrite generation in aerated cultures. Because CRP is a transcription factor, we reasoned it might play a role in oxygen-responsive transcriptional activation of the *napA*, *napB*, and *napC* genes as well as the gene encoding the nitrate-responsive histidine kinase NarQ (49). Therefore, in an aerated Δ*crp* mutant culture, transcription of these genes would not increase in response to nitrate as they would in aerated WT and *ΔhapR* mutant cultures. However, we found that the *nap* genes and *narQ* were similarly regulated regardless of genetic background and aeration. (Fig 2D and Fig S2). We conclude that CRP does not activate transcription of the *narQ* and *nap* genes in shaking cultures supplemented with nitrate and that another function of CRP must enhance nitrite generation in aerated cultures.

### Evidence that conversion of nitrite to nitric oxide is not responsible for inhibition of biofilm accumulation by nitrite

Nitric oxide is known to cause dispersal of the *V. cholerae* biofilm (50, 51). However, the *V. cholerae* genome does not encode the enzymes that convert nitrite to nitric oxide, and spontaneous reduction of nitrite to nitric oxide seems unlikely in the oxidizing conditions of aerobic cultures (51). Nevertheless, to test this possibility, we added the nitric oxide scavenger 2- carboxyphenyl-4,4,5,5-tetramethylimidazoline-1-oxyl-3-oxide (c-PTIO) to *V. cholerae* cultured statically in LB supplemented with nitrite (51). As shown in Fig S3, this treatment did not increase biofilm accumulation in nitrite-supplemented cultures.

### CRP is essential for nitrate reduction by a Δ*hapR* mutant in aerated cultures and for the resistance of the Δ*hapR* mutant biofilm to inhibition by nitrate and nitrite

To further probe the resistance of the Δ*hapR* mutant biofilm to inhibition by nitrate and nitrite, we created a Δ*hapR*Δ*crp* double mutant and tested its ability to convert nitrate to nitrite and to form biofilms in the presence of nitrate under static and aerated conditions. Similar to the Δ*crp* mutant, the double mutant was unable to convert of nitrate to nitrite in aerated cultures (Fig 2C). In static cultures, biofilm accumulation by the Δ*hapR*Δ*crp* double mutant in the presence of nitrate and nitrite was negligible and total growth was similar to that of the Δ*crp* mutant (Fig 3A). Therefore, deletion of *crp* in the Δ*hapR* mutant background restored the susceptibility of the Δ*hapR* mutant biofilm to inhibition by nitrate and nitrite. While these findings do not elucidate the mechanism underlying the resistance of the Δ*hapR* mutant to nitrate and nitrite, they do allow us to predict the contribution of *V. cholerae* VPS-dependent biofilm formation to surface colonization under conditions that might be encountered in the mammalian intestine provided the following two assumptions are true. (i) In a glucose-rich environment, CRP is inactive and WT *V. cholerae* responds to nitrate as a Δ*crp* mutant would. (ii) At low cell density, HapR is inactive, and WT *V. cholerae* responds to nitrate as a Δ*hapR* mutant would. If this is correct, the following model shown in Fig 3B should be valid. In glucose-rich media, *V. cholerae* reduction of nitrate to nitrite should occur in static but not aerated cultures. In glucose-rich environments, this leads to inhibition of biofilm formation by nitrate in static but not aerated cultures independent of cell density (Fig 3B (pathways Ia and b and Ila and b)). In environments such as the terminal ileum, where glucose is limiting, *V. cholerae* reduces nitrate to nitrite regardless of aeration, but only high cell density cultures are susceptible to inhibition by nitrite (Fig 3B (pathways Ic and llc).

**Figure 3:**
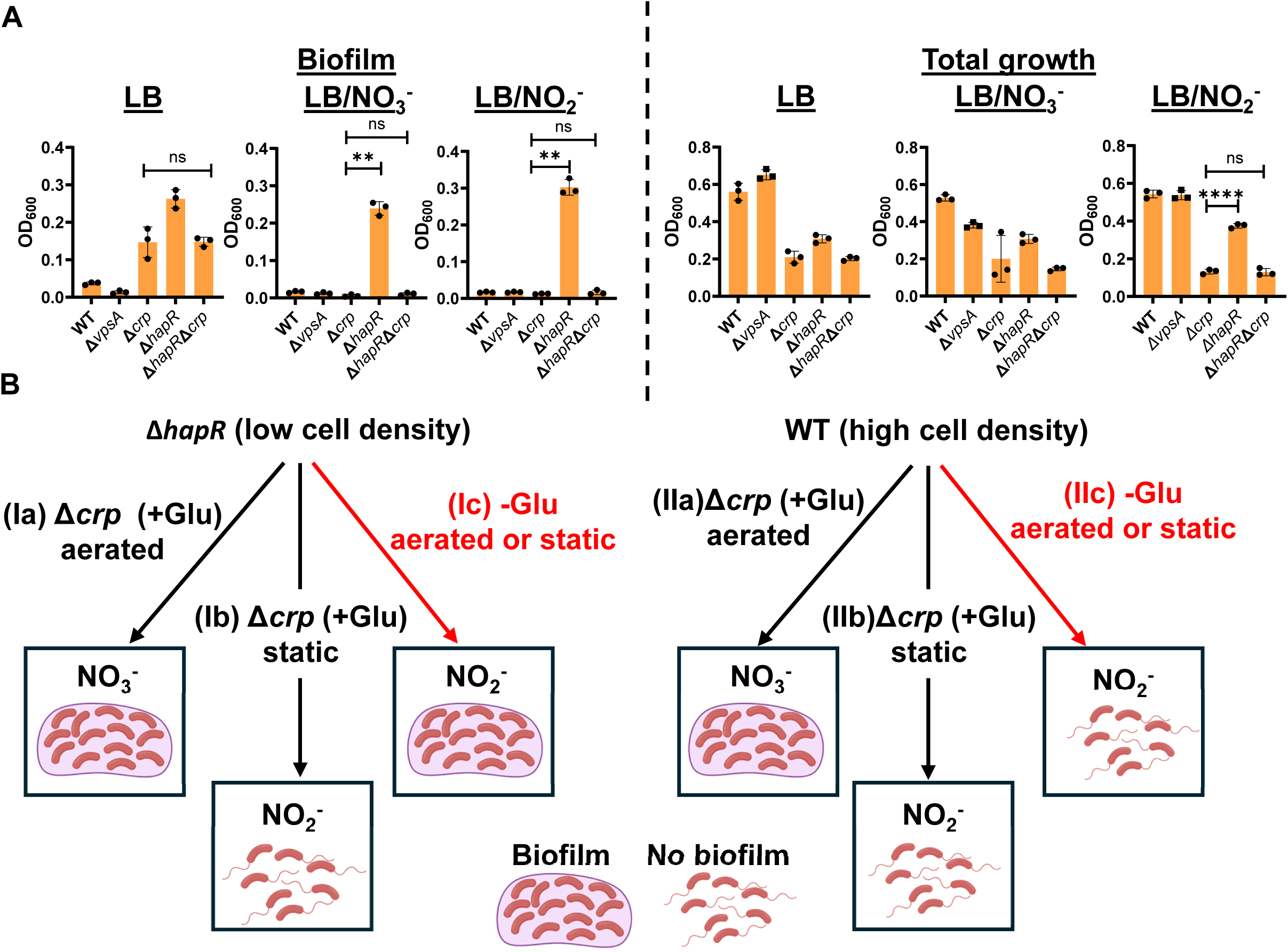
CRP and HapR govern *V. cholerae* biofilm inhibition by nitrate. (A) Quantification of biofilm accumulation and total growth by WT *V. cholerae* as well as Δ*crp*, Δ*hapR*, and Δ*hapR*Δ*crp* mutants cultured statically in LB alone or supplemented with 5 mM nitrate (NO_3_^-^) or nitrite (NO_3_^-^). The mean of biological triplicates is shown for all experiments. Error bars reflect the standard deviation. For biofilm accumulation, significance was calculated using a Brown-Forsythe and Welch’s ANOVA with a Dunnett’s T3 multiple comparisons test. For total growth, significance was calculated using an ordinary one-way ANOVA with a Dunnett’s multiple comparisons test. **** p<0.0001, *** p<0.001, ** p<0.01, * p<0.05, ns not significant. (B) A model showing the impact of carbon source, cell density, and aeration on inhibition of *V. cholerae* biofilm formation by nitrate. This model is based on our observations using the Δ*crp* mutant, which mimics *V. cholerae* behavior in a glucose-rich environment (+Glu), and the Δ*hapR* mutant, which mimics cells in a low cell density state. (Ia and lla) In a Δ*crp* mutant background or when glucose is plentiful and cultures are aerated [Δ*crp* (Glu+), aerated], nitrate (NO_3_^-^) conversion to nitrite (NO_2_^-^) is blocked in both Δ*crp* (equivalent to glucose-rich, high cell density) and Δ*hapR*Δ*crp* (equivalent to glucose-rich, low cell density) mutants, and biofilm accumulation proceeds. (lb and IIb) In a Δ*crp* mutant or when glucose is plentiful and cells are cultured statically [Δ*crp* (Glu+), static], nitrate conversion to nitrite proceeds in both Δ*crp* (equivalent to glucose-rich, high cell density) and Δ*hapR*Δ*crp* (equivalent to glucose-rich, low cell density) mutants, and biofilm accumulation is inhibited. (Ic and IIc) When glucose is scarce (-Glu), nitrate is converted to nitrite in both WT *V. cholerae* (high cell density) and the Δ*hapR* mutant (low cell density). The Δ*hapR* mutant (low cell density) biofilm is resistant to nitrite, while the WT biofilm (high cell density) is inhibited by nitrite. These pathways are highlighted in red because they are the only ones in which nitrate regulation of biofilms differs at high and low cell density. (Created in BioRender. Watnick, P. (2025) https://BioRender.com/n69b437)

### Nitrate has a global impact on the *V. cholerae* transcriptome

We reasoned that genes involved in inhibition of *V. cholerae* biofilm accumulation by nitrate should be differentially regulated in response to nitrate in WT *V. cholerae* and the Δ*crp* and Δ*hapR*Δ*crp* mutants but not the Δ*hapR* mutant. To gain insight into the genetic basis of *V. cholerae* biofilm inhibition by nitrate, we carried out an RNAseq experiment including WT *V. cholerae* and Δ*crp*, Δ*hapR*, and Δ*hapR*Δ*crp* mutants cultured statically for 8 hours in LB alone or supplemented with 5 mM nitrate (Tables S1-4). In general, more RNAs were differentially regulated in response to nitrate in the Δ*hapR* mutant than in the other genetic backgrounds tested. The Δ*crp* mutant had the fewest differentially regulated RNAs, and the Δ*hapR*Δ*crp* mutant had an intermediate number (Fig 4A). This suggests that CRP plays an important role in the transcriptional and post-transcriptional response of *V. cholerae* to nitrate, while HapR mutes the transcriptional response to nitrate. Only 74 coding sequences, 22 intergenic RNAs, and 14 antisense RNAs were commonly differentially regulated in the 4 strains in response to nitrate (Fig 4B). Gene ontology (GO) analysis of coding sequences using the Panther Overrepresentation test showed significant overrepresentation of nitrate-regulated genes involved in aerobic respiration and iron transport but not surface adhesion (Fig 4C and Tables S1-4) (52, 53). Genes involved in iron transport were overrepresented in all genetic backgrounds. One possible explanation is that the ferric uptake regulator Fur, which represses transcription of iron uptake genes in response to ferrous iron availability, is less active in the presence of nitrate (Fig 4C) (54). We can conclude that the differential regulation of iron transport genes in response to nitrate is not responsible for the distinct responses of the Δ*crp* and Δ*hapR* mutant to nitrate.

**Figure 4:**
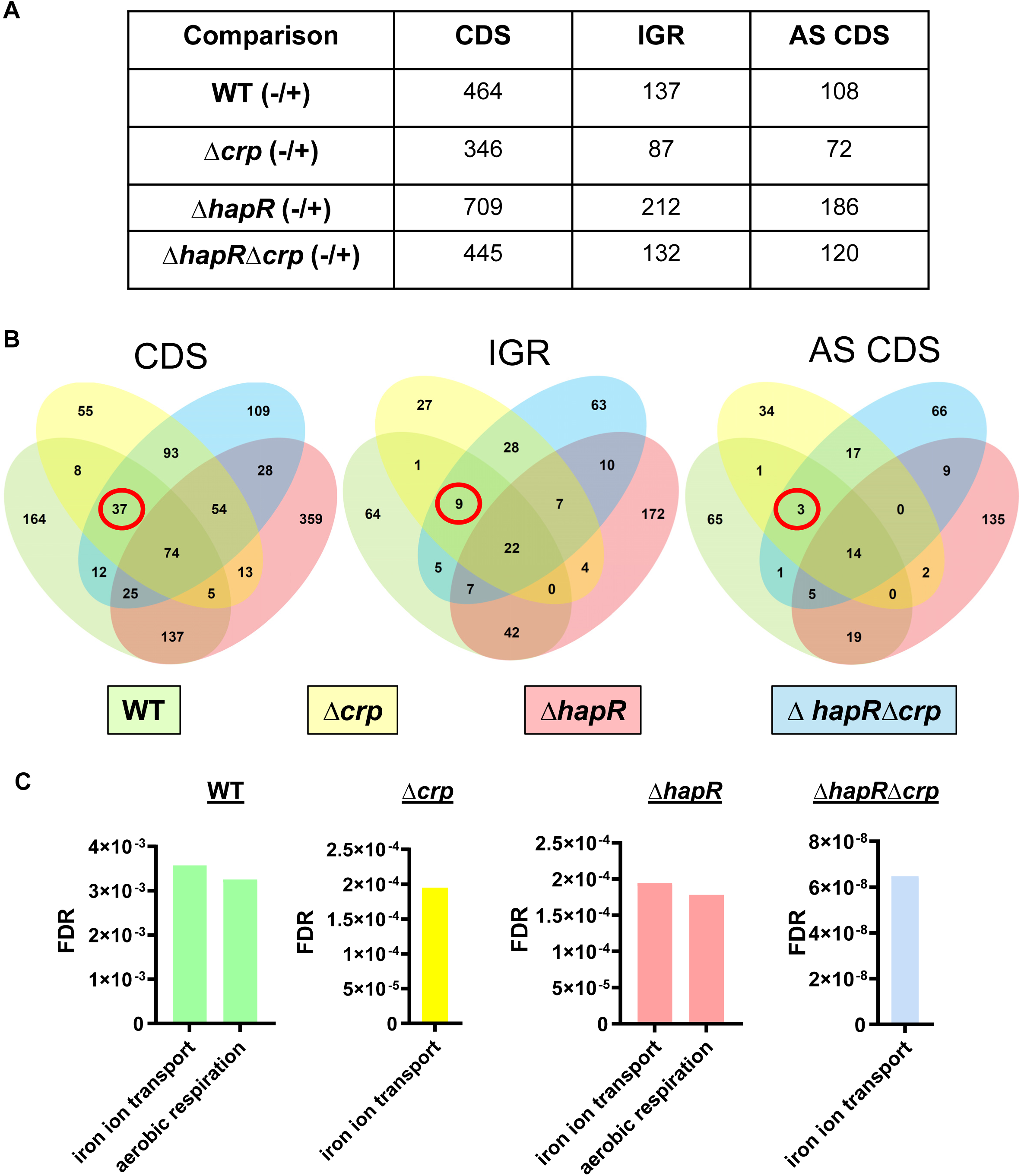
Nitrate regulation of aerobic respiration is CRP-dependent, while nitrate control of iron-regulated genes is CRP and HapR-independent. (A) Table showing numbers of differentially regulated coding sequences (CDS), intergenic region RNAs (IGR), and antisense RNAs within coding sequences (AS) for the indicated strains when cultured statically for 8 hours either alone (-) or in the presence of 5 mM nitrate (+). (B) Venn diagrams illustrating overlap in transcripts differentially regulated by nitrate in each of the genetic backgrounds studied. Transcripts found in coding sequences (CDS), intergenic regions (IGR), and antisense RNA (AS CDS) are illustrated separately. Genes that are differentially regulated in all but the Δ*hapR* mutant are circled in red. (C) Gene ontology analysis of differentially regulated genes using the Panther Overrepresentation test.

Surprisingly, nitrate supplementation activated not only transcription of the genes required for nitrate respiration, but also those required for respiration of oxygen. However, the latter were only differentially regulated by nitrate when CRP was present (Fig 4C). Indeed, transcription of *bd-l* and *cbb3*, the genes encoding the principal aerobic terminal oxidases of *V. cholerae*, was greatly reduced in the Δ*crp* mutant in both the presence and absence of nitrate (Fig S4) (23). We have previously shown that oxygen inhibits nitrate reduction. Based on these findings, we hypothesize that, due to a paucity of aerobic terminal oxidases, the Δ*crp* mutant cannot efficiently deplete aerated cultures of oxygen, and this may underlie the inability of the Δ*crp* mutant to generate nitrate.

### Nitrate does not decrease *V. cholerae* biofilm accumulation via known pathways

With the exception of Bap1 (VC1888), the genes responsible for forming the *V. cholerae* biofilm matrix are found in two islands, *vps-*I, which includes VC0916-27, and *vps-*II, which includes VC0934-9 (3, 6, 43, 55, 56). These genes are regulated by two transcription factors, VpsR (VC0665) and VpsT (VCA0952), both of which are activated by binding of the second messenger cyclic-diguanylate (c-di-GMP) (57–61). We first scrutinized transcription of these genes (Fig 5A). Differential regulation of most of these genes did not meet our criteria for fold-change and statistical significance. Genes that were significantly regulated did not follow the pattern expected based on biofilm phenotype. For instance, while nitrate decreased WT *V. cholerae* biofilm accumulation, transcription of biofilm genes was increased in response to nitrate. Conversely, while nitrate had no impact on the Δ*hapR* mutant biofilm, transcription of biofilm genes was decreased in response to nitrate. To confirm these puzzling patterns of gene regulation, we undertook qRT-PCR analysis for a subset of these genes (Fig 5B). Supporting our RNAseq findings, nitrate activated the biofilm gene *vpsA* in WT *V. cholerae* and repressed the biofilm genes *vpsR*, *vpsT*, and *vpsA* in the Δ*hapR* mutant. These results suggest that nitrate does not inhibit *V. cholerae* biofilm accumulation by decreasing transcription of the biofilm synthesis genes.

**Figure 5:**
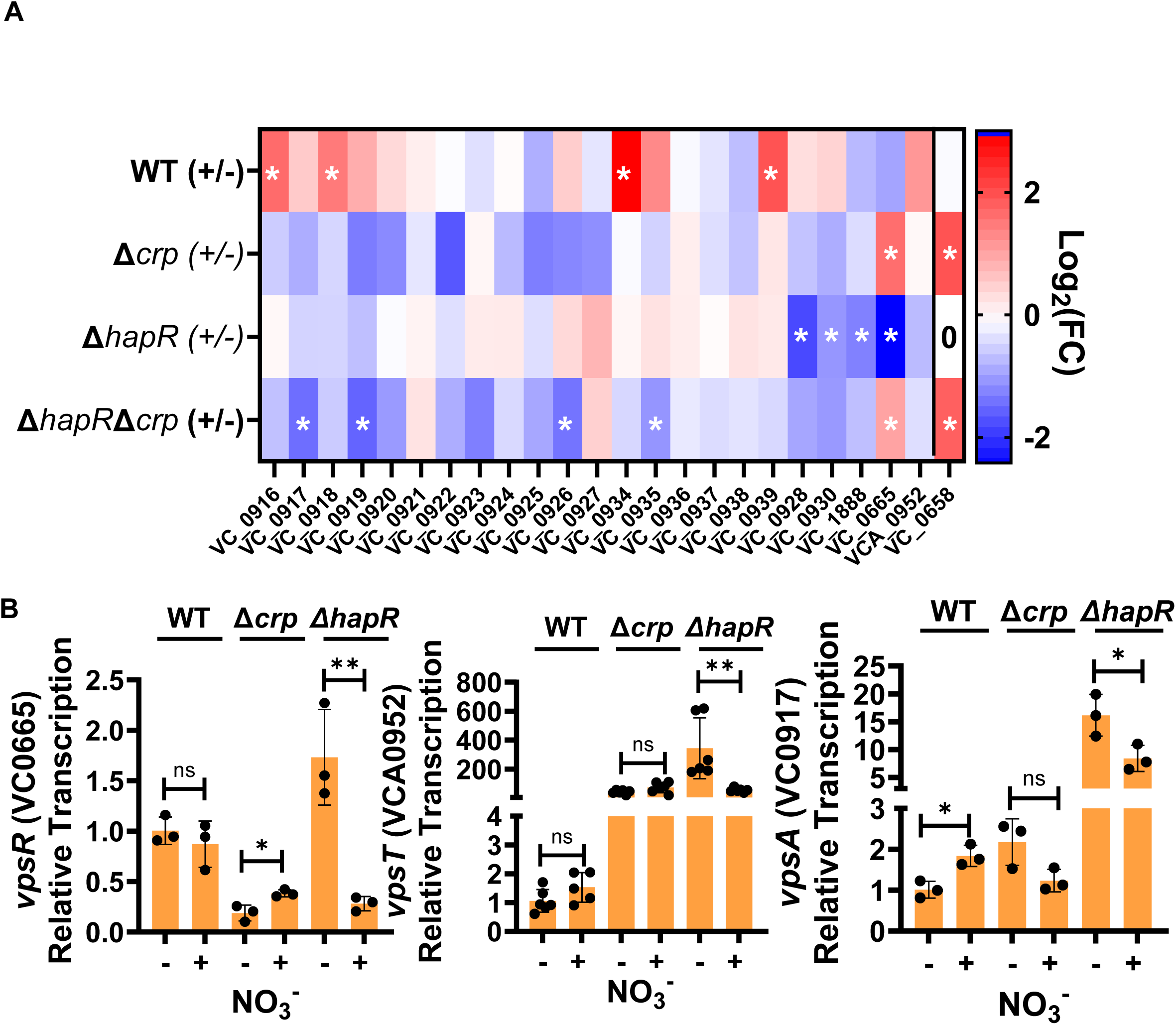
Repression of the *vps* genes is not responsible for inhibition of *V. cholerae* biofilm accumulation by nitrate. (A) Heat map illustrating differential regulation of biofilm formation and biofilm dispersal genes as measured by RNAseq. Fold-change represents transcription in +nitrate/- nitrate conditions. White stars indicate genes that met the criteria for significance FC>2 and padj < 0.05.(B) qRT-PCR confirmation of *vps* gene transcription in the presence (+) and absence (-) of nitrate. The indicated strains were cultured statically for 8 hours. The mean of biological triplicates is shown. Error bars reflect the standard deviation. Significance was calculated using a student’s t test. ** p<0.01, * p<0.05, ns not significant.

We then explored known post-transcriptional mechanisms of biofilm regulation. Two previously identified sRNAs, *ryhB* and *qrrX*, regulate *V. cholerae* biofilm formation. The Fur-regulated small RNA *ryhB*, whose transcription was increased with nitrate supplementation for all genetic backgrounds including the Δ*hapR* mutant, activates biofilm formation (62). Since *ryhB* activates biofilms, it is unlikely that *ryhB* is responsible for biofilm inhibition by nitrate (63). *qrrX* is an RNA sponge that destabilizes the *qrr1-4* sRNAs, which maintain low cell density signaling and transcription of the *vps* genes (64, 65). *qrrX* transcription increased in the presence of nitrate in WT *V. cholerae*, which would be expected to decrease biofilm formation, but transcription of *qrrX* is quite low in Δ*crp*, Δ*hapR*, and Δ*crp*Δ*hapR* mutants (Fig. S5). Therefore, *qrrX* also seems an unlikely candidate for biofilm inhibition. There were many additional coding and putative non-coding RNAs within this dataset that have not previously been associated with biofilm inhibition and, therefore, warrant further exploration.

*V. cholerae* accumulates on surfaces at low cell density, and HapR contributes to biofilm dispersion at high cell density (48). Therefore, we entertained the alternative hypothesis that nitrate induces biofilm dispersal at high cell density rather than inhibiting biofilm matrix synthesis. Several manuscripts have reported genes that are essential for *V. cholerae* biofilm dispersal (66–69). We reasoned that transcription of biofilm dispersal genes might be activated in response to nitrate in all genetic backgrounds tested except Δ*hap*R, which mimics a low cell density state. A total of 49 genes and non-coding RNAs were differentially regulated in all but the Δ*hapR* mutant background (Fig 4B, red circle). However, this group did not include any genes known to be involved in biofilm formation or dispersal. Examining genes that were differentially regulated in the Δ*crp* and Δ*hapRΔcrp* mutant backgrounds only and unchanged in WT *V. cholerae* and the Δ*hapR* mutant, we identified one gene VC0658 or *cdgI* previously identified to be involved in biofilm dispersal (Fig 5A) (66). CdgI is a c-di- GMP phosphodiesterase and, therefore, would be predicted to decrease intracellular levels of c-di- GMP and thereby *vps* gene transcription. Because we did not observe a decrease in *vps* gene transcription in response to nitrate, it seems unlikely that CdgI is responsible for biofilm inhibition by nitrate unless it is acting via a novel, c-di-GMP-independent mechanism. We conclude that, if HapR regulates the biofilm response to nitrate by increasing biofilm dispersal, this is unlikely to occur through a previously described pathway.

### *Paracoccus aminovorans*, a microbe associated with susceptibility to cholera, rescues *V. cholerae* biofilm accumulation in the presence of nitrate

In a human host, inhibition of *V. cholerae* VPS-dependent biofilm formation due to nitrate reduction to nitrite might diminish colonization of the ileum, where the concentration of nitrate is high. We reasoned that a nitrite- reducing microbial community might mitigate inhibition of VPS-dependent biofilm formation and, thus, predispose to *V. cholerae* colonization and disease. To test this, we co-cultured *V. cholerae* with the aerobic Gram-negative rod *Paracoccus aminovorans*. Sequenced strains of *P. aminovorans* JCM 7685 and DSM 8537 (NCBI Bioprojects PRJEB8789 and PRJEB17339) encode the genes for reduction of nitrite to nitric oxide, nitrous oxide, and finally nitrogen gas. Furthermore, in a prospective study of household contacts of cholera patients, the presence of *P. aminovorans* in stool was associated with subsequent development of disease (8). In a later publication, *P. aminovorans* was found to increase colonization of abiotic surfaces and the neonatal mouse intestine via formation of a VPS-dependent biofilm (7). Using the *P. aminovorans* strain previously isolated from cholera stool and generously provided by the Weil and Ng labs, we first cultured *P. aminovorans* statically in LB supplemented with 5 mM nitrate or 5 mM nitrite and quantified nitrite remaining after 48 hours. As shown in Figs 6 A and B, nitrite was not present in the medium after 48 hours of culture, suggesting that *P. aminovorans* is able to reduce nitrite. To determine whether *P. aminovorans* might reduce the nitrite generated from nitrate by *V. cholerae*, we statically cultured WT *V. cholerae* and the Δ*crp* mutant alone or with *P. aminovorans* in LB supplemented with nitrate and measured the nitrite concentration after 48 hours. In both co-cultures, *P. aminovorans* greatly diminished nitrite concentrations in the medium at the end of the incubation period (Fig 6B). To investigate whether reduction of nitrite by *P. aminovorans* might rescue *V. cholerae* surface attachment, we compared *V. cholerae* surface adhesion in LB alone or supplemented with nitrate in the presence and absence of *P. aminovorans.* As shown in Fig. 6C and D, the presence of *P. aminovorans* rescued biofilm accumulation by WT and Δ*crp* mutant *V. cholerae* in the presence of nitrate. Interestingly, *P. aminovorans* also significantly improved the growth of the Δ*crp* mutant, suggesting it secretes an as yet unidentified metabolite that can be utilized by Δ*crp* mutant. *P. aminovorans* made a very little biofilm on its own and did not increase surface adhesion by a Δ*vps* mutant. This demonstrates that *P. aminovorans* rescues VPS-dependent biofilm formation in the presence of nitrate rather than providing *V. cholerae* an alternate means of surface adherence. To determine the contribution of *V. cholerae* and *P. aminovorans* to the biofilm mass measured, we formed biofilms with *V. cholerae*, which is streptomycin (Sm)-resistant, and a Sm-sensitive *P. aminovorans* strain. After 48 hours, biofilms were dispersed and colony forming units (CFU) were enumerated on LB agar alone or supplemented with Sm. As shown in Fig S6, CFU were comparable regardless of the addition of antibiotics, suggesting that *V. cholerae* is the major contributor to measured biofilm biomass in co-cultures. We conclude that members of the microbiota that reduce nitrite can rescue *V. cholerae* biofilm accumulation in the presence of nitrate.

**Figure 6:**
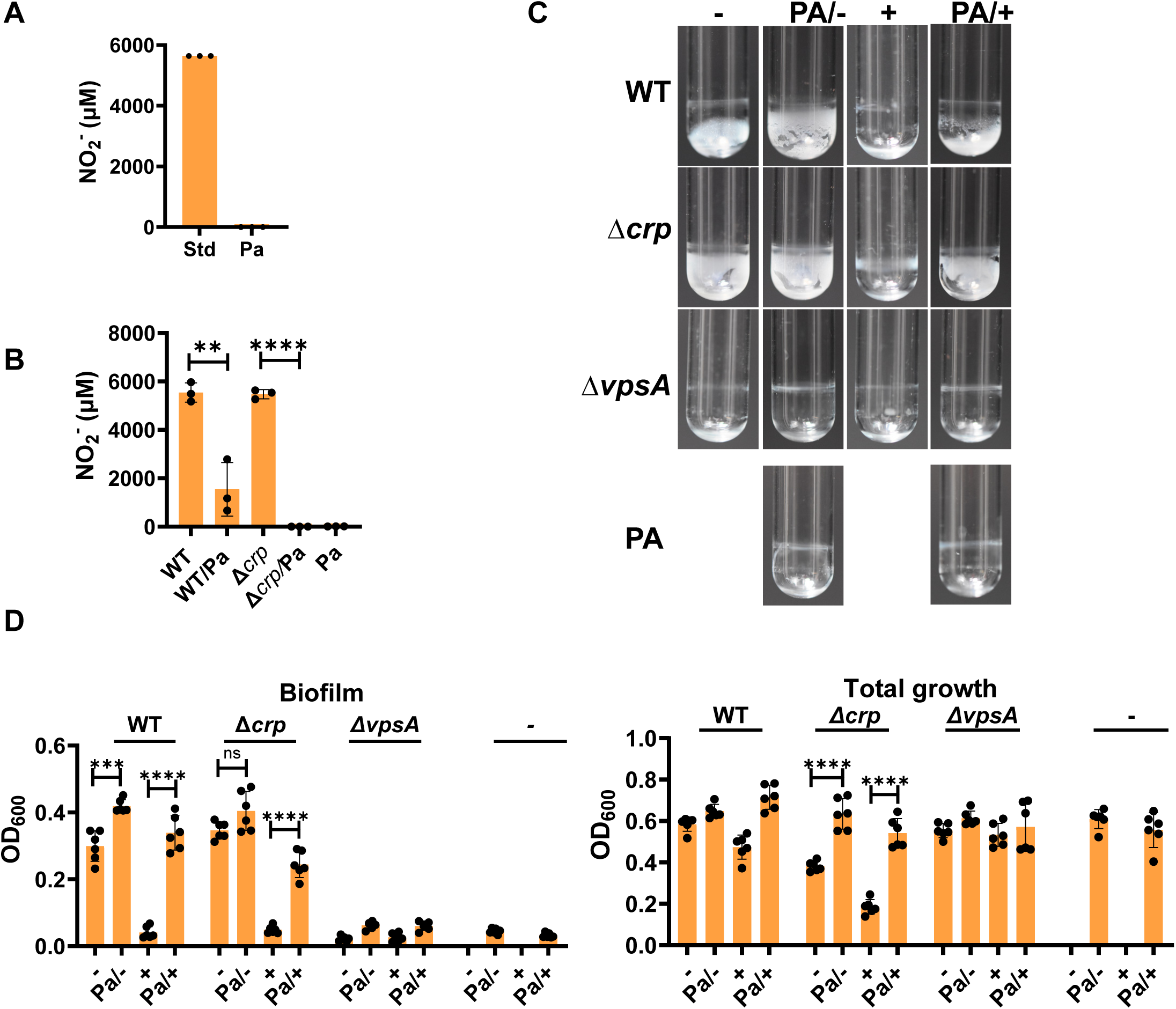
*Paracoccus aminovorans* consumes nitrite produced by *V. cholerae* and partially rescues *V. cholerae* biofilm accumulation in the presence of nitrate. (A) Quantification of nitrite in sterile LB broth supplemented with 5 mM nitrite (Std) and the spent supernatant of a *P. aminovorans* (Pa) cultured statically for 48 hours in LB supplemented with 5 mM nitrite (Pa). (B) Quantification of nitrite in the supernatants of WT *V. cholerae* and the indicated mutants cultured statically for 48 hours alone or with *P. aminovorans* (Pa) in LB supplemented with 5 mM nitrate. The mean of biological triplicates is shown. Error bars reflect the standard deviation. Significance was calculated using a student’s t test. (C) Representative images of biofilms formed by WT *V. cholerae* and the indicated mutants cultured statically for 48 hours alone or with *P. aminovorans* (Pa) in LB (-) or LB supplemented with 5 mM NaNO_3_ (+). (D) Quantification of biofilm formation and total growth by the indicated strains cultured as described in (C). The mean of six biological replicates is shown. Error bars reflect the standard deviation. For biofilm accumulation, significance was calculated using a Welch’s t test. For total growth, significance was calculated using a student’s t test. **** p<0.0001, ***p <0.001, ** p<0.01, * p<0.05, ns not significant.

## Discussion

Here we report that *V. cholerae* vigorously respires nitrate in shaking cultures to produce nitrite, that CRP, a transcription factor that is active in the absence of glucose, is essential for aerobic reduction of nitrate to nitrite, and that nitrite inhibits *V. cholerae* biofilm accumulation when HapR is active at high cell density but not in a Δ*hapR* mutant, which mimics low density behavior. Based on our findings, we propose that inhibition of *V. cholerae* biofilm accumulation by nitrate is controlled by carbon source and cell density. Finally, we show that co-culture with *P. aminovorans*, an inhabitant of the human intestinal microbiota that increases susceptibility to cholera, respires nitrite and augments *V. cholerae* surface accumulation in the presence of nitrate. These findings are significant because they highlight the role of the microbiota in regulation of surface colonization by *V. cholerae* in environments such as the mammalian terminal ileum where both oxygen and nitrate are available as terminal electron acceptors. They also leave unanswered questions that are discussed below.

We have not elucidated the mechanism by which CRP activates nitrate reduction in aerated cultures although we have ruled out transcriptional activation of the *narQ and nap* genes as the basis for this. Our RNAseq data suggest that expression of the principal terminal oxidases bd-I and cbb3 is decreased in a Δ*crp* mutant leading to impaired oxygen respiration. Previous work suggests that aerobic bacterial cultures progress towards a hypoxic state as cell density increases. We predict that this would occur more slowly in a Δ*crp* mutant (70). Our data show that aeration delays nitrate reduction by wild-type *V. cholerae*. If this is a direct response to oxygen, nitrate reduction should be further delayed in a Δ*crp* mutant. Two conserved and well-studied transcription factors respond to oxygen availability and could contribute to the inability of the Δ*crp* mutant to reduce nitrate in aerated cultures. The first is the fumarate nitrate reduction regulator FNR, which is active as a dimer when its (4Fe-4S)^2+^ cluster cofactor is bound. In the presence of O2, FNR dimerization is inhibited by sulfur oxidation of this co-factor (71). Therefore, FNR is a direct sensor of oxygen availability. In a previous study, FNR was identified as essential for nitrate reduction by *V. cholerae* in anaerobic cultures, but transcriptional targets of FNR were not further explored (30). In other organisms, FNR regulates additional genes found to be essential for *V. cholerae* nitrate reduction including the *moa* genes, encoding the molybdenum cofactor biosynthesis proteins, and the *nqr* genes encoding the NADH:ubiquinone oxidoreductase (72–74). The ArcAB two component system also controls gene transcription in response to oxygen availability (75). ArcB is a membrane-associated sensor that responds to anaerobiosis by activating the transcription factor ArcA via phosphorylation. ArcB is thought to respond oxygen availability by sensing the oxidation state of quinones that shuttle electrons in aerobic electron transport chains (76). In *V. cholerae*, ArcA activates biofilm formation via *vpsT*, represses flagellar synthesis, and increases expression of the virulence factors cholera toxin and the toxin co-regulated pilus (77–79). Because this regulation occurs at the transcriptional level, it does not explain the observed response to nitrate. In other organisms, both FNR and ArcAB also control expression of small RNAs (sRNAs) that regulate protein expression (27, 80–82). In our RNAseq experiment, many putative sRNAs were differentially transcribed in response to nitrate, and it is quite likely that one or more of these contributes to regulation of nitrate reduction at the post-transcriptional level.

We also have not uncovered the mechanism by which nitrite inhibits *V. cholerae* VPS-dependent biofilm formation nor the basis for the resistance of the Δ*hapR* mutant biofilm to nitrite. Our experiments rule out transcriptional regulation of the *vps* genes as a mechanism, and it seems unlikely that nitrite would directly poison the enzymes involved in WT and Δ*crp* biofilm formation but not that of the Δ*hapR* mutant, since biofilm formation by all depends on the same suite of enzymes. While HapR has previously been associated with activation of biofilm dispersal at high cell density, we have not identified a plausible role for known biofilm dissolution genes in HapR-dependent inhibition of *V. cholerae* biofilm accumulation by nitrite (48). Our RNAseq experiments suggest that HapR regulates many putative sRNAs in response to nitrate. We hypothesize that one of these is responsible for post-transcriptional inhibition of biofilm formation in the presence of nitrite.

Finally, we have not explored the basis of increased transcription of iron scavenging genes in nitrate- supplemented *V. cholerae* cultures. These genes are regulated by the ferric uptake repressor Fur. Fur represses transcription of iron transport genes by binding reduced or ferrous iron (Fe^2+^) (83). Under the conditions of our RNAseq experiment, while we expect most of the available iron to be in the ferric or oxidized state (Fe^3+^), there is sufficient Fe^2+^ to promote repression of transcription by Fur (84, 85). One possibility is that addition of nitrate decreases the availability of Fe^2+^, but this cannot be explained by differential regulation of *V. cholerae vciB* (VC0283), which is essential for iron reduction in the periplasm, as this gene was not differentially regulated in the presence of nitrate (Tables S1-4) (86, 87). Furthermore, while NapC has been shown to participate in iron reduction, its contribution is relatively small (87, 88). Therefore, competition between Fe^3+^ and nitrate for reduction by NapC is not a likely mechanism. We have discussed possible roles for FNR and ArcAB in the response to nitrate. Highlighting the complexity of the bacterial response to electron acceptor availability, in other organisms, molybdenum cofactor biosynthesis and nitrate metabolism is under control of Fur, FNR, and ArcA, and ArcA regulates both Fur and uptake of ferrous iron (89–92). A methodical examination of this complex regulatory network will be required to understand activation of iron-regulated genes by nitrate.

Our studies highlight key unexplored aspects of *V. cholerae* surface association in nitrate-rich environments that are likely to have implications for disease. *V. cholerae* colonization of the ileum, a region of the intestine where oxygen is present and nitrate concentrations reach 6 mM, is critical for expression of *V. cholerae* virulence factors (25). Previous studies have shown that CRP is essential for robust *V. cholerae* colonization of the mammalian intestine (93). We propose that in the glucose- poor environment of the ileum, *V. cholerae* CRP is active, available oxygen is consumed, and nitrate is reduced (Fig 7). Thus, *V. cholerae* colonizes the ileum in the presence of nitrite until high density is reached. HapR then activates a response to nitrite that inhibits or disperses the biofilm. Because HapR also represses expression of the toxin co-regulated pilus, cholera toxin, and the biofilm synthesis genes, we propose that HapR enacts a regulatory program that limits *V. cholerae* colonization of the intestine and prolongs survival of the host (94–96). Despite the actions of HapR, some hosts are differentially susceptible to cholera (9). We suggest that intestinal microbes such as *P. aminovorans*, which convert the nitrite generated by *V. cholerae* nitrate respiration to inactive byproducts, short-circuit HapR-induced biofilm inhibition in response to nitrate.

**Figure 7:**
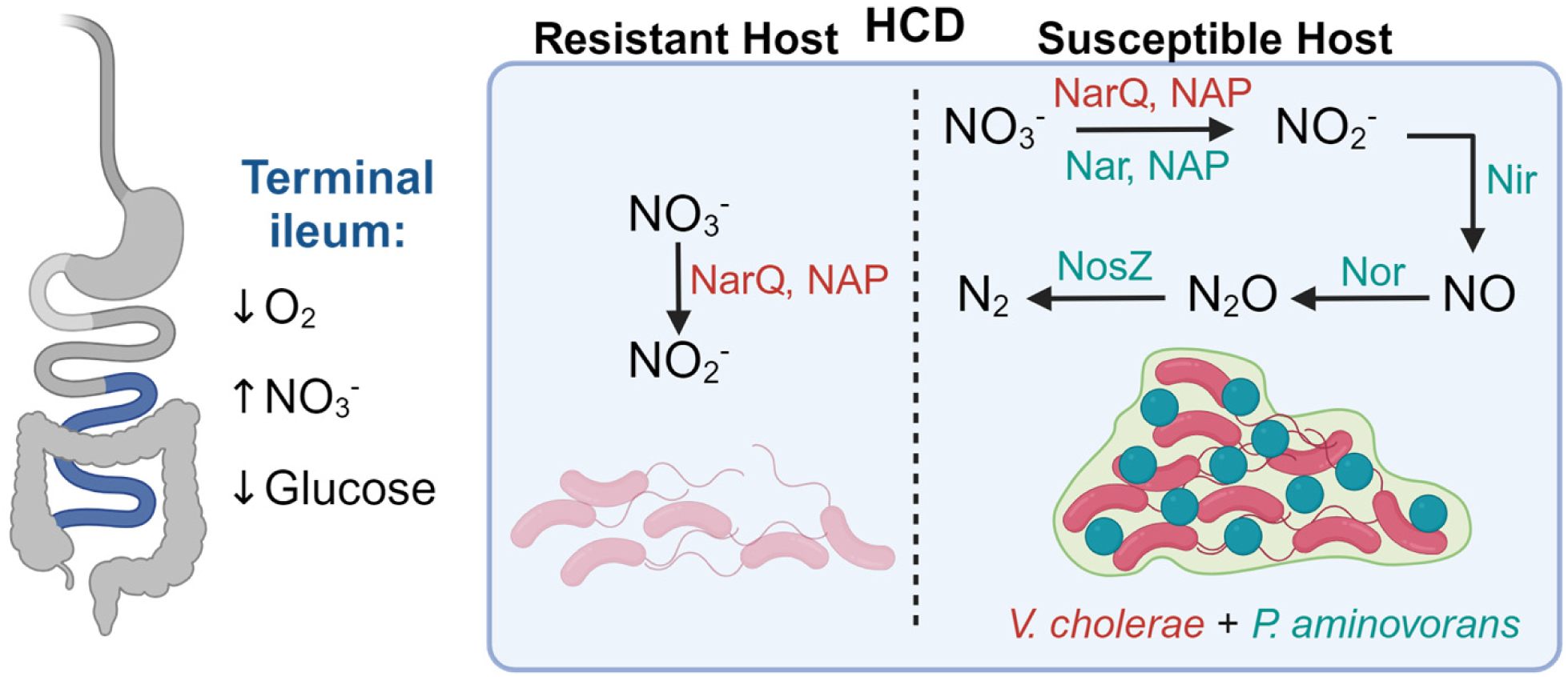
A model for *V. cholerae* high cell density colonization of the ileum in the presence and absence of a nitrite-reducing microbe such as *P. aminovorans*. Schematic showing the impact of nitrite-reducing organisms such as *P. aminovorans* on *V. cholerae* intestinal colonization at high cell density (HCD) and host susceptibility to cholera. A microbial community metagenome that encodes the *nir*, *nor*, and *nosZ* genes, which further reduce nitrite to nitric oxide (NO), nitrous oxide (N2O), and nitrogen gas (N2), respectively, may enhance *V. cholerae* surface accumulation at high cell density and contribute to increased susceptibility to cholera. (Created in BioRender. Watnick, P. (2024) BioRender.com/c85w494 )

## Materials and Methods

### Bacterial Strains and Media

The quorum sensing-competent *V. cholerae* strain C6706, which is Sm-resistant, served as the parental strain for all experiments. Sm-sensitive and Sm-resistant *P. aminovorans* strains were generously provided by Drs. Wai-Leung Ng and Ana Weil. The Sm-sensitive strain was previously isolated from stool swabs of a family member of a cholera patient in Bangladesh (7, 8). Both *V. cholerae* and *P. aminovorans* strains were cultured with shaking in LB broth (Difco, Miller formulation) supplemented with Sm (100 µg/mL, Gibco) unless otherwise noted. *Escherichia coli* strains used for cloning were cultured in LB broth supplemented with ampicillin (100 µg/mL, Sigma). For rescue experiments, 0.5 mM isopropyl-β-D-thiogalactopyranoside (IPTG, MedChemExpress) was used to induce expression. *V. cholerae* was cultured at 27 °C, while *P. aminovorans* and *E. coli* were cultured at 37°C. Except for the Δ*napC* mutant, all plasmids and strains used were previously published and are shown in Table S5. The Δ*napC* mutant was generated by double homologous recombination as previously described (33). The following primers were used to generate the deletion construct, which was cloned into pWM91(97): primer 1: AATTGGGTACCGGGCCCCCCCCTGGCTGGCATTGCTCAT, primer 2: TCCTTATTGCCAGCCTTCTGGGCTCGATAGACGAAG, primer 3: GAAGGCTGGCAATAAGGA, primer 4: CTTATCGATACCGTCGACCTCGAGCAGGAAACCACCATGAG.

### Biofilm Assays

Cells were cultured overnight, diluted 1:100 into fresh LB broth and cultured to mid-exponential phase using the conditions described above. Mid-exponential cells were diluted in the indicated medium to yield an OD_600_ of 0.05 as measured on a SpectraMax ABS microplate reader (Molecular Devices). For *V. cholerae* biofilm assays, 300µl of this diluted culture was aliquoted into 10x75mm borosilicate glass tubes (Fisher) and incubated for approximately 20 h at 27 °C either statically or with agitation at 200 rpm as noted. Tubes were positioned vertically for static cultures, and at an angle for aerobic cultures to increase exposed surface area. A similar procedure was followed for co-cultures with *P. aminovorans* with the following variations. Where noted, a 1:1 ratio of *P. aminovorans* to *V. cholerae* was diluted into a 300 μl volume in a 10x75mm borosilicate tube and cultured statically at room temperature for 48 hours. Planktonic cells were removed and replaced with an equal volume of phosphate-buffered saline as well as several 1 mm glass beads. Biofilm-associated cells were then dispersed by vortexing. The density of biofilm and planktonic cells was quantified by measuring an OD_600_. Total growth was calculated as the sum of the planktonic and biofilm OD_600_. A minimum of three biological replicates was performed, and each experiment was repeated at least once with similar results. Biofilms were imaged using a Nikon D3400 digital camera.

### Nitrite measurements

Overnight cultures were diluted 1:100 in LB medium supplemented with 5 mM sodium nitrate or nitrite as noted and cultured at 27 °C for the indicated time. Prior to the measurements, cultures were centrifuged at 12,000 x g for 1 min at room temperature. The supernatants were collected and filtered through a 0.2 µm filter. A kit based on the Griess reagent (Biotium) was used to measure nitrite concentrations per the manufacturer’s instructions. Briefly, 50 µL of the Griess reagent was combined with 150 µL of the sample and 1.3 mL of ddH2O and incubated at RT for at least 30 minutes prior to measurement of A548 using an SpectraMax iD5 microplate reader (Molecular Devices). The concentration of nitrite was determined by plotting the experimental values against a standard curve generated from a 1 mM stock of nitrite solution. Biological triplicates were performed, and each experiment was repeated at least once with similar results.

### RNA isolation

Total RNA was isolated from 3 mL cultures cultivated at 27°C without agitation for 8 hours. Bacterial pellets were collected and resuspended in 500 µl TriZol (Invitrogen), incubated at 60°C for 10 min, followed by purification using Direct-zol RNA Miniprep kit (Zymo R2052) following manufacturer’s instructions. RNA for both RNAseq analysis and qRT-PCR was prepared using this method.

### Generation of RNA-Seq data

Illumina cDNA libraries were generated using a modified version of the RNAtag-seq protocol (98). Briefly, 0.5-1 μg of total RNA was fragmented, depleted of genomic DNA, dephosphorylated, and ligated to DNA adapters carrying 5’-AN8-3’ barcodes of known sequence with a 5’ phosphate and a 3’ blocking group. Barcoded RNAs were pooled and depleted of rRNA using the riboPOOLS rRNA depletion kit (siTOOLs Biotech). Pools of barcoded RNAs were converted to Illumina cDNA libraries in 2 main steps: (i) reverse transcription of the RNA using a primer designed to the constant region of the barcoded adaptor with addition of an adapter to the 3’ end of the cDNA by template switching using SMARTScribe (Clontech), and (ii) PCR amplification using primers whose 5’ ends target the constant regions of the 3’ or 5’ adaptors and whose 3’ ends contain the full Illumina P5 or P7 sequences. cDNA libraries were sequenced on the Illumina NovaSeq 6000 platform to generate paired end reads.

### RNAseq data analysis

Sequencing reads from each sample in a pool were demultiplexed based on their associated barcode sequence using custom scripts. Up to 1 mismatch in the barcode was allowed provided it did not make assignment of the read to a different barcode possible. Barcode sequences were removed from the first read as were terminal G’s from the second read that may have been added by SMARTScribe during template switching. Reads were aligned to *Vibrio cholerae* N16961 genome, and read counts were assigned to genes and other genomic features using custom scripts. Differential expression analysis was conducted with DESeq2 (99) and EdgeR (100). Genes were considered significantly differentially regulated if a two-fold or greater change in expression was measured and if the adjusted p value was less than or equal to 0.05. RNAseq data were visualized using the IGV (v2.10.3) developed at the Broad Institute. Genome ontology analysis utilized Panther version 19.0 (52, 101). Significantly overrepresented biological functions were identified using a Fisher exact test for significance and p<0.05.

### qRT-PCR

500 ng of total RNA served as a template to make c-DNA using the iScript reverse transcriptase supermix (Bio-Rad). A QuantStudio 5 Real-Time PCR machine was used to measure amplification of cDNA added to an iTaq Universal SYBR Green Supermix reaction (Bio-Rad). *rpoA* was used as a reference gene for static cultures, while *clpX* was used for aerated cultures. Technical duplicates and biological triplicates were performed.

### Data analysis

Data were graphed and analyzed for statistical significance using Prism (Graphpad). A mean and standard deviation were calculated. For qRT-PCR, nitrite generation, and total growth measurements, a student’s t test or one way ANOVA was used to evaluate significance. For biofilm accumulation measurements, a Welch’s t test or Brown-Forsythe and Welch’s ANOVA was used to evaluate significance.

## Data availability statement

RNAseq data have been deposited in the NCBI repository (accession no. GSE272711).

## Acknowledgements.

This work was funded by NIH AI112652 to P.I.W. The *Paracoccus aminovorans* strain was generously provided by Drs. Ana Weil and Wai-Leung Ng. RNA-Seq libraries were constructed and sequenced at the Broad Institute of MIT and Harvard by the Microbial ‘Omics Core and Genomics Platform, respectively. The Microbial‘Omics Core also provided guidance on experimental design and conducted preliminary analysis for all RNA-Seq data.

---

**Table S1: RNAseq analysis of WT *V. cholerae* cultured statically in the absence and presence of 5 mM NaNO_3_ (-/+).**

**Table S2: RNAseq analysis of a *V. cholerae* Δ*crp* mutant cultured statically in the absence and presence of 5 mM NaNO_3_ (-/+).**

**Table S3: RNAseq analysis of a *V. cholerae* Δ*hapR* mutant cultured statically in the absence and presence of 5 mM NaNO_3_ (-/+).**

**Table S4: RNAseq analysis of a *V. cholerae* Δ*hapR*Δ*crp* mutant cultured statically in the absence and presence of 5 mM NaNO_3_ (-/+).**

**Table S5: Strains, plasmids, and primers used in this work.**

## Supplemental Materials

**Figure S1:**
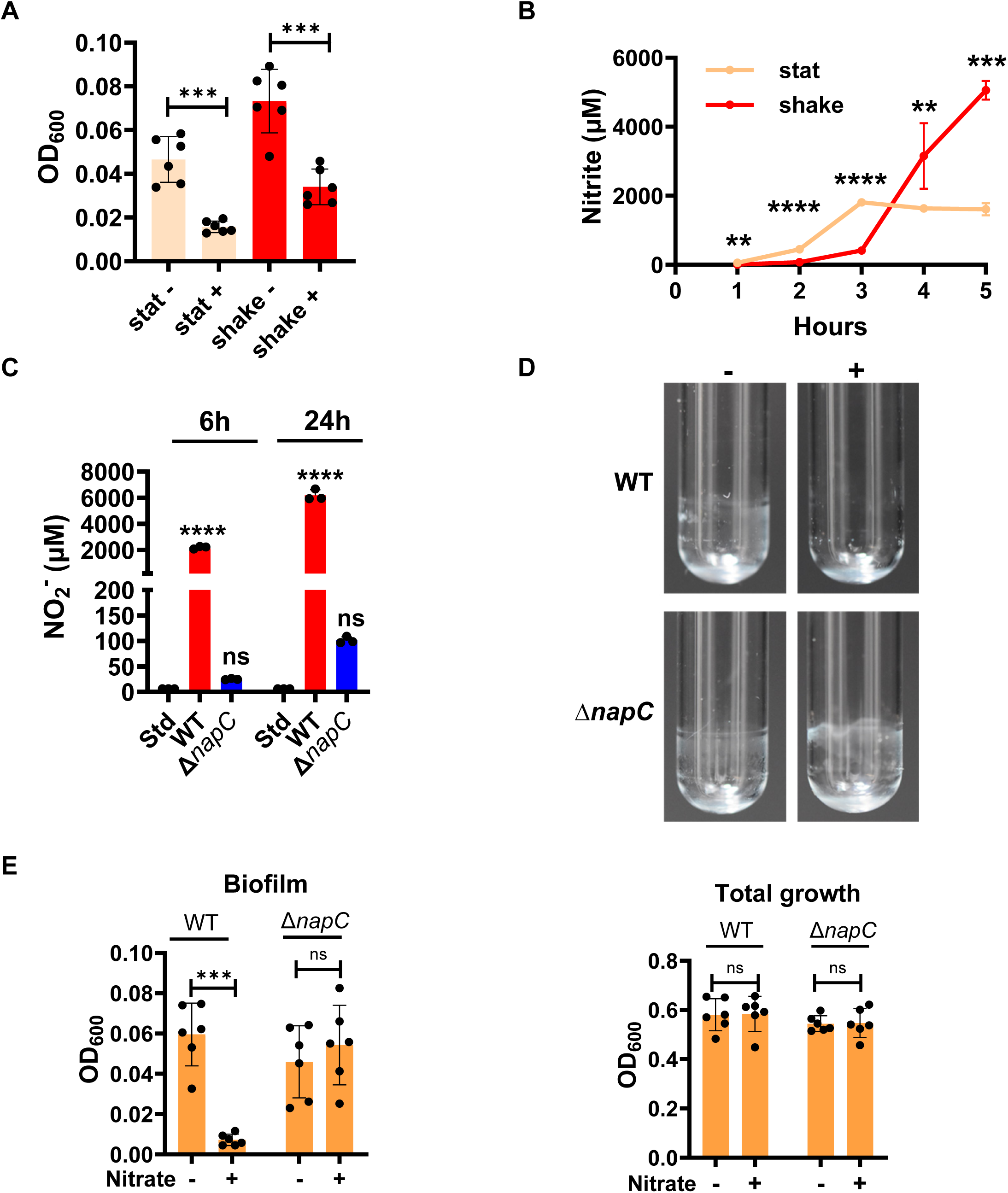
Nitrate reduction to nitrite is essential for inhibition of biofilm accumulation by nitrate. (A) An expanded view of the WT *V. cholerae* biofilm measurement shown in Fig 1D. (B) Nitrite generation by WT *V. cholerae* under static and aerated conditions. The mean of biological triplicates is shown. Error bars represent the standard deviation. Significance at each time point was evaluated using a Welch’s t test. (C) Quantification of nitrite (NO_2_^-^) generation in LB supplemented with nitrate by WT *V. cholerae* and a Δ*napC* mutant after 6 and 24 hours of incubation. Std indicates a 5 mM nitrate in LB standard without bacteria added. The mean of biological triplicates is shown. Error bars represent the standard deviation. An ordinary one-way ANOVA with Dunnett’s test was used to determine whether measurements differed significantly from the standard. (D) Images of biofilms formed by WT *V. cholerae* and a Δ*napC* mutant in LB alone (-) or supplemented with 5 mM nitrate (+). (E) Quantification of biofilm formation and total growth by the strains in (D). The mean of six biological replicates is shown. Error bars represent the standard deviation. Significance was calculated using a Welch’s t test for biofilm accumulation and a student’s t test for total growth. ****p<0.0001, ***p<0.001, ** p<0.01, ns not significant.

**Figure S2:**
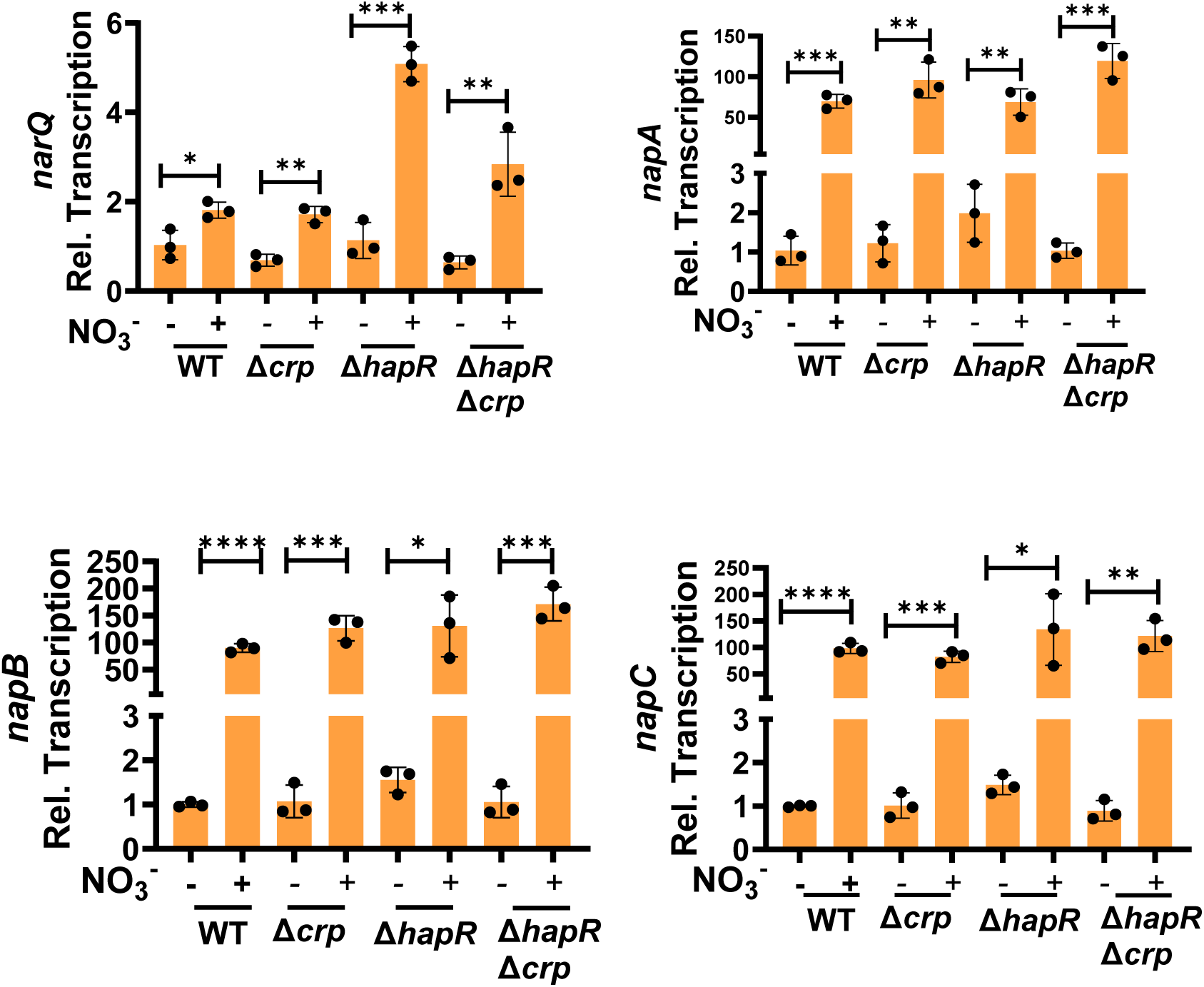
Expression of the *narQ* and the *nap* genes increases in response to nitrate supplementation in static cultures of WT *V. cholerae* and mutants. qRT-PCR quantification of transcription of the *V. cholerae* genes required for nitrite generation in LB alone (-) and supplemented with 5 mM nitrate (+). The indicated strains were cultured statically and harvested after 8 hours. The mean of biological triplicates is shown for all experiments. Error bars reflect the standard deviation. Significance was calculated using a student’s t test. **** p<0.0001, *** p<0.001, ** p<0.01, * p<0.05, ns not significant.

**Figure S3:**
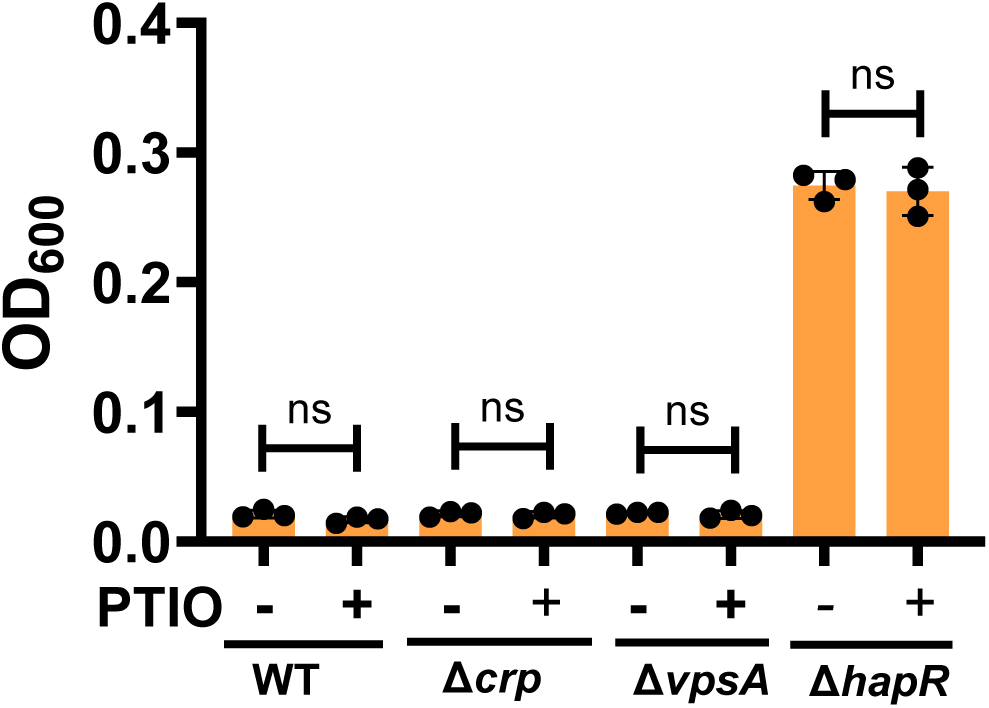
The nitric oxide scavenger PTIO does not rescue inhibition of *V. cholerae* biofilm formation by nitrite. Quantification of biofilms formed by WT *V. cholerae* and the indicated mutants cultured statically in LB supplemented with 5 mM nitrite alone (-) or with addition of the nitric oxide scavenger 2-phenyl- 4,4,5,5-tetramethylimidazoline-1-oxyl-3-oxide (PTIO, 500 µM) (+). The mean of biological triplicates is shown. Error bars reflect the standard deviation. Significance was calculated using a student’s t test. ns not significant.

**Figure S4:**
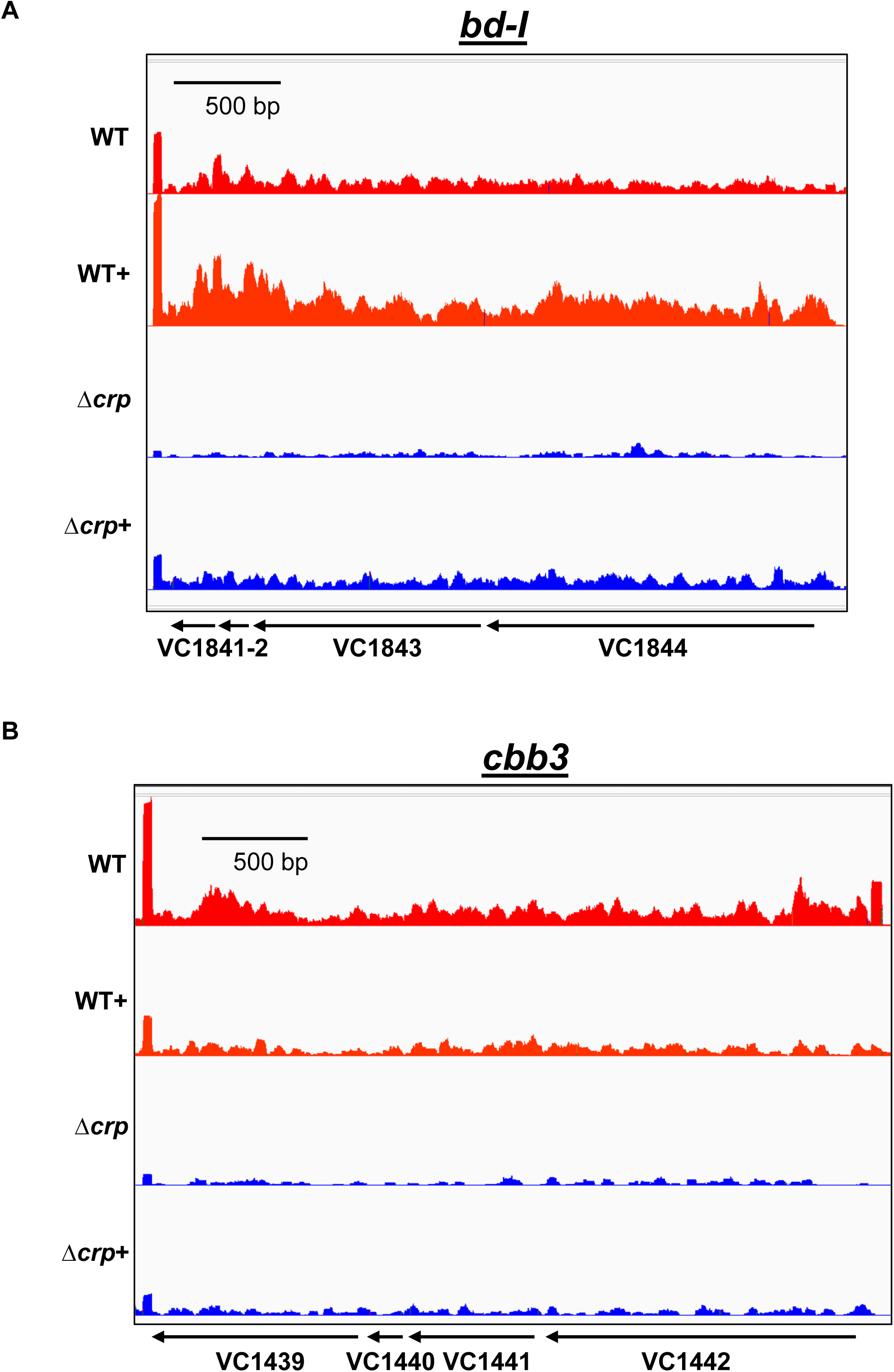
CRP activates transcription of the principal aerobic terminal oxidases of *V. cholerae*. Normalized RNAseq data showing transcription of the principal aerobic terminal oxidases of *V. cholerae* (A) *bd-I* and (B) *cbb3* in WT *V. cholerae* and the Δ*crp* mutant. Cells were cultured statically in Lb broth alone (-) or supplemented with 5 mM nitrate (+). Experiments were performed in triplicate. A representative trace is shown.

**Figure S5:**
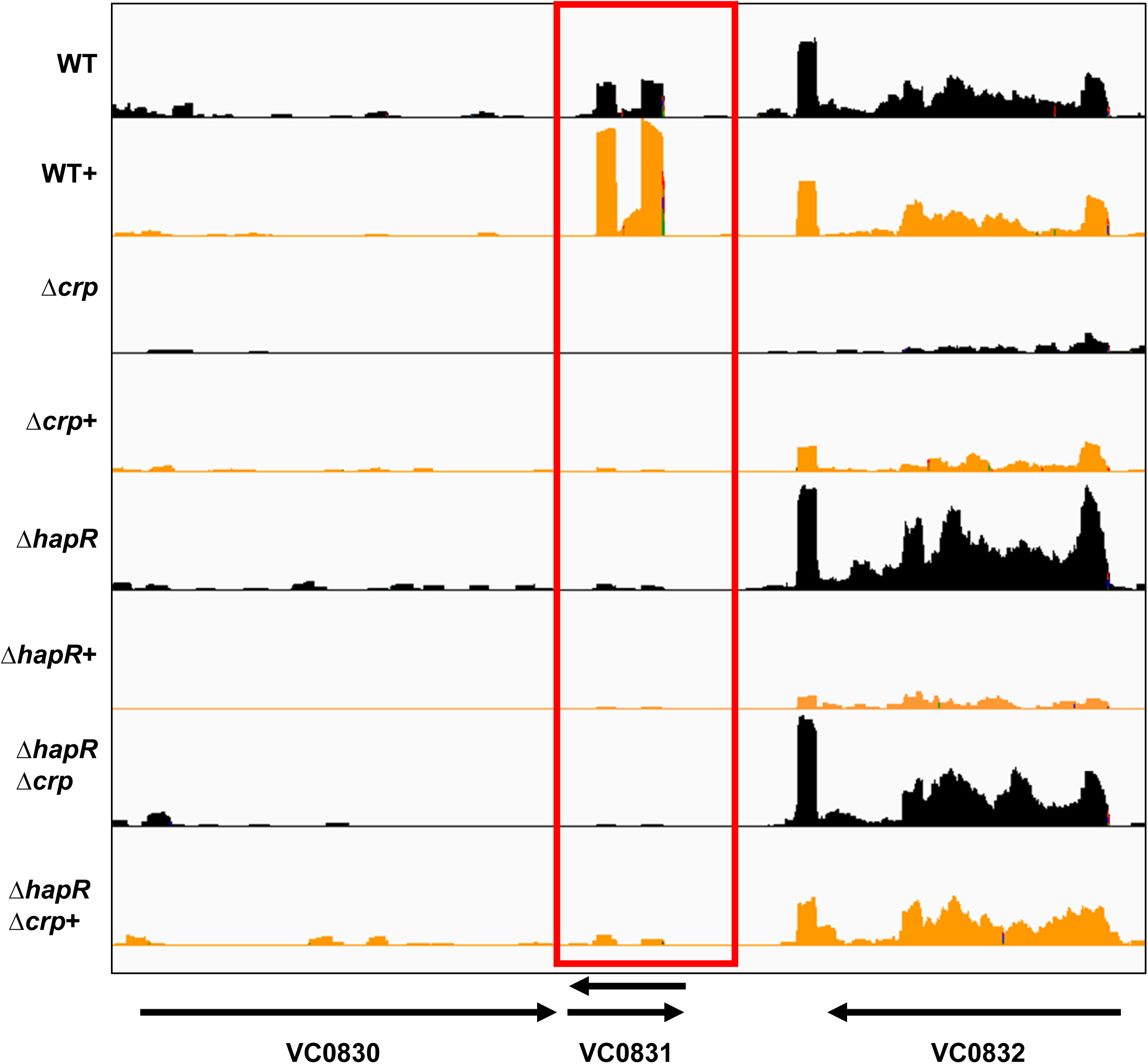
*qrrX* is regulated by nitrate but does not follow the pattern expected of a nitrate-responsive biofilm inhibitor. Normalized RNAseq data showing expression of *qrrX*, which is located between VC0830 and VC0832 and highlighted by the red box. Biological triplicates were performed. A representative trace is shown here.

**Figure S6:**
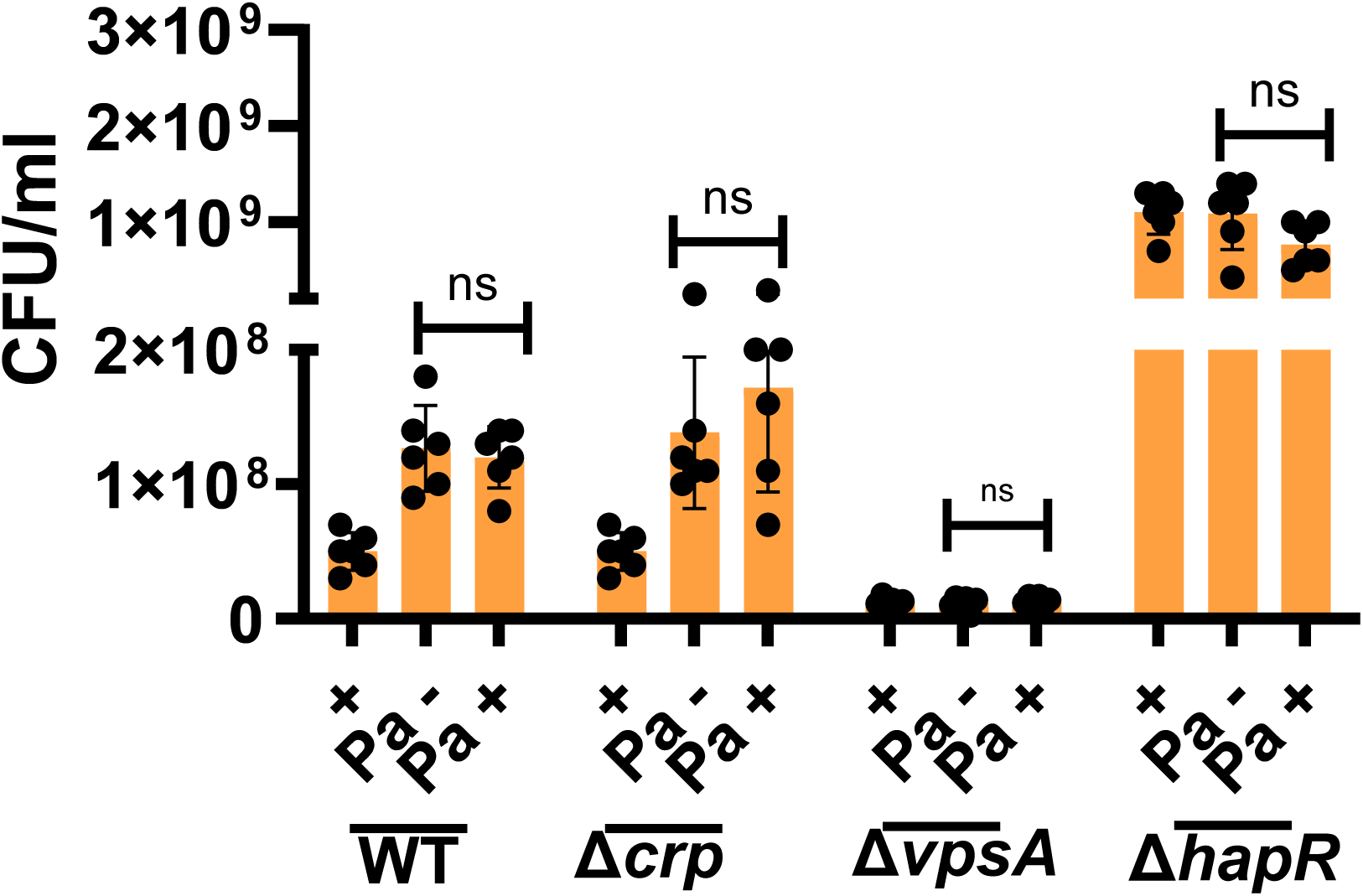
The biofilm formed when *Paracoccus aminovorans* and *V. cholerae* are co-cultured in the presence of nitrate consists principally of *V. cholerae* cells. Quantification of total and streptomycin-resistant (Sm) colony forming units (CFU/ml) in biofilms formed by WT *V. cholerae* and the indicated mutants, which are all streptomycin-resistant, alone or in co-culture with streptomycin-sensitive *P. aminovorans* (Pa). Cells were cultured statically for 48 hours in medium supplemented with 5 mM nitrate alone (-) or with added streptomycin (+). Biofilm mass was quantified by dispersal in phosphate buffered saline, plating of serial dilutions, and enumeration of CFU. The mean of six biological replicates is shown. Error bars reflect the standard deviation. Significance was calculated using a student’s t test. ns not significant.

